# High dose IFN-*β* activates GAF to enhance expression of ISGF3 target genes in epithelial cells

**DOI:** 10.1101/2020.10.28.358366

**Authors:** Kensei Kishimoto, Catera Wilder, Justin Buchanan, Minh Nguyen, Chidera Okeke, Alexander Hoffmann, Quen J. Cheng

## Abstract

Interferon *β* (IFN-*β*) signaling activates the transcription factor complex ISGF3 to induce gene expression programs critical for antiviral defense and host immune responses. It has also been observed that IFN-*β* activates a second transcription factor, γ-activated factor (GAF), but the significance of this coordinated activation is unclear. We report that in respiratory epithelial cells high doses of IFN-*β* indeed activate both ISGF3 and GAF, which bind to distinct genomic locations defined by their respective DNA sequence motifs. In contrast, low doses of IFN-*β* preferentially activate ISGF3 but not GAF. Surprisingly, in epithelial cells GAF binding does not induce nearby gene expression even when strongly bound to the promoter. Yet expression of interferon stimulated genes is enhanced when GAF and ISGF3 are both active compared to ISGF3 alone. Our data suggest that GAF enhances ISGF3 target gene expression by co-localizing with ISGF3 at some promoters and facilitating chromosome looping between distal enhancers and other promoters. We propose that GAF may function as a dose-sensitive amplifier of ISG expression to enhance antiviral immunity and establish pro-inflammatory states in respiratory epithelial cells.

**One sentence summary:** GAF is transcriptionally inactive in epithelial cells but enhances expression of ISGF3 target genes, thus functioning as a dose-sensitive amplifier of the IFN-*β* response.

## Introduction

The type I interferon (IFN) response is essential to host antiviral immunity. Respiratory epithelial cells, particularly alveolar type II cells, are the primary target of respiratory pathogens and are capable of both producing and responding to IFNs (*1–7*). Type I IFN signaling induces expression of IFN stimulated genes (ISGs), which code for antiviral effector molecules as well as chemokines, cytokines, and other immune effector molecules (*5, 8*). Consequently, defects in type I IFN signaling result in susceptibility to various infections, while overexuberant IFN responses are associated with heightened and pathological inflammation (*9–14*). Therefore, the regulation of type I IFN signaling is critically important, as its mis-regulation in either direction can lead to disease.

Type I IFNs signal through the type I IFN receptor (IFNAR) composed of IFNAR1 and IFNAR2 subunits (*15*). IFNAR is associated with the Janus tyrosine kinases JAK1 and TYK2. Upon ligand binding, the IFNAR bound JAK1/TYK2 kinases cross-phosphorylate and activate signal transducer and activator of transcription (STAT) proteins STAT1 and STAT2. Phosphorylated STAT1 and STAT2 interact with IRF9 to form the trimeric transcription factor (TF) ISGF3, which binds to interferon-stimulated response elements (ISREs) defined by the sequence AGTTTCn_2_TTTC (*16*).

STAT1 may also form a homodimer (*17*) named *γ*-activated factor (GAF), which is typically induced by stimulation with type II IFN, IFN-*γ* (*18*). GAF binds to gamma-activated sequences (GAS), defined by the motif TTCn_2-4_GAA (*19*), and IFN-*γ* has a well-described role in immune cells such as macrophages (*20*). Notably, however, type I IFN signaling can also activate GAF in both immune and non-immune cells (*1, 18, 21, 22*). Although this has been observed for decades, the function of IFN-β-induced GAF is unclear. It is possible that GAF and ISGF3 have redundant roles in gene regulation, as type I and type II IFNs were reported to induce highly overlapping gene expression programs (*23, 24*). It has also been suggested that a primary role of IFN-*β*-induced GAF is to induce the secondary TF IRF1, which can increase basal ISG expression (*25*) or amplify expression of pro-inflammatory genes such as *Gpb5* (*22*). To date, however, few studies have systematically examined the role of IFN-*β*-induced GAF using genome-wide approaches.

Here we report that in murine epithelial cells IFN-β activates both ISGF3 and GAF but in differing amounts depending on IFN-β dose. We demonstrate that at high-doses of IFN-β, GAF enhances the ISGF3-responsive expression of ISGs genomewide, not through the direct transcriptional activity of GAF but by a variety of genespecific indirect mechanisms.

## Materials and Methods

### Cell culture

MLE-12 cells obtained from ATCC (Manassas, VA) were cultured in DMEM (Corning Inc., Corning, NY) supplemented with 10% FBS, 100 IU/ml penicillin, 100 μg/ml streptomycin, 2 mM L-glutamine at 37^0^C at 5% CO_2_. Recombinant murine IFN-β (PBL Assay Science, Piscataway, NJ) and IFN-γ (BD Biosciences, San Jose, CA) were used for stimulation.

### ChlP-seq libraries

MLE-12 cells grown to 80% confluence in 15-cm plates were double crosslinked with 2 mM disuccinimidyl glutarate in PBS for 30 minutes and 1% methanol-free formaldehyde in PBS for 10 minutes at room temperature (RT). Crosslinking reactions were quenched with 125 mM of glycine for 5 minutes at RT, followed by washing of the cells twice with cold PBS supplemented with 1 mM EDTA. Crosslinked cells were then scraped in PBS with EDTA, collected by centrifugation, and frozen at −80 °C. Cell pellets were thawed in Lysis Buffer 1 (50 mM HEPES-KOH pH 7.6, 140 mM NaCl, 1 mM EDTA, 10% glycerol, 0.5% NP-40, 0.25% Triton X-100, and 1x protease inhibitor cocktail (Thermo Scientific, Waltham, MA)) on ice and incubated at 4 °C. for 7 min. Cells were then sonicated with Diagenode Bioruptor 300 sonication system with medium intensity and 15 seconds ON / 30 seconds OFF for 3 cycles in 1.5mL TPX microtubes and centrifuged to isolate nuclei. Nuclei were washed in Lysis Buffer 2 (10 mM Tris-HCl pH 8.0, 200 mM NaCl, 1 mM EDTA, 0.5 mM EGTA, and 1x protease inhibitor cocktail) at RT for 10 mins. After centrifugation, nuclei were resuspended in Lysis Buffer 3 (10 mM Tris-HCl pH 8.0, 100 mM NaCl, 1 mM EDTA, 0.5 mM EGTA, 0.1% Na Deoxycholate, 0.5% N-lauroylsarcosine, and 1x protease inhibitor cocktail) and sonicated in TPX microtubes at low intensity for 30 sec ON / 30 sec OFF for a total of 12 cycles while inverting and pulse-spinning after every 4 cycles. After centrifuging the samples at max speed for 10 mins at 4 °C., supernatant containing fragmented chromatin was diluted with 6 volumes of Dilution Buffer (10 mM Tris-HCl pH 8.0, 160 mM NaCl, 1 mM EDTA, 0.01% SDS, 1.2% Triton X-100, 1x protease inhibitor cocktail) and incubated with Protein-G Dynabeads (Thermo Fisher) for 2 hours at 4 °C for pre-clearing. The chromatin was then incubated with 5 mg of STAT1 antibody (Cell Signaling Technologies, Danvers, MA; CST-9172) for every ½ plate of cells at 4 °C overnight.

The antibody-chromatin complexes were then immunoprecipitated by incubating with protein-G Dynabeads for 5 hours at 4 °C. Beads were collected and washed twice with the following buffers: Low Salt buffer (50 mM HEPES-KOH pH 7.6, 140 mM NaCl, 1 mM EDTA, 1% Triton X-100, 0.1% Na Deoxycholate, 0.1% SDS), High Salt buffer (50 mM HEPES-KOH pH 7.6, 500 mM NaCl, 1 mM EDTA, 1% Triton X-100, 0.1% Na Deoxycholate, 0.1% SDS), LiCl buffer (20 mM Tris-HCl pH 8.0, 250 mM LiCl, 1 mM EDTA, 0.5% Na Deoxycholate, 0.5% NP-40), and TE buffer (10 mM Tris-HCl pH 8.0, 1 mM EDTA). Immunoprecipitated chromatin complexes were treated with RNAse A (Thermo Fisher) at 37 °C for one hour, and crosslinks were reversed with 10% SDS and 0.6 mg/ml Proteinase K (New England Biolabs, Ipswich, MA) overnight with shaking at 65 °C. Immunoprecipitated DNA fragments were purified with AMPure XP SPRI beads (Beckman Coulter, Brea, CA) at a 0.95 volume ratio according to manufacturer’s instructions.

DNA was quantified with Qubit dsDNA HS (Life Technologies, Carlsbad, CA) using a Qubit 2.0 fluorometer. Libraries were prepared for sequencing using NEBNext Ultra II DNA Library Prep Kit (New England Biolabs) with NEBNext Multiplex Oligos (New England Biolabs). Each library was prepared from 1-5 ng of starting DNA. Input samples were pooled from samples from the same experiment. Final libraries were checked for quality by agarose gel, quantified with Qubit, and multiplexed with a maximum of 24 samples per sequencing reaction. Libraries were sequenced on an Illumina HiSeq 2500 with single end 50bp reads at the UCLA Broad Stem Cell Research Center.

### RNA-seq libraries

Total RNA was purified from Trizol with DIRECTzol kit (Zymo Research, Irvine, CA) according to manufacturer’s instructions. For data in Fig. 3b, Fig. 4g, Fig. 5, Fig. S2, and Fig. S3, RNA-seq libraries were prepared using KAPA stranded mRNA-seq library kit per manufacturer’s instructions and single-end sequenced at read length 50 bp on an Illumina HiSeq 2500. For data in Fig. 3e–3d, Fig. 4a, and Fig. 4c-f, RNA-seq libraries were generated by MedGenome, Inc. (Foster City, CA) using Illumina stranded TruSeq mRNA kits and paired-end sequenced at read length 150 bp on an Illumina NovaSeq 6000.

**Figure 1:**
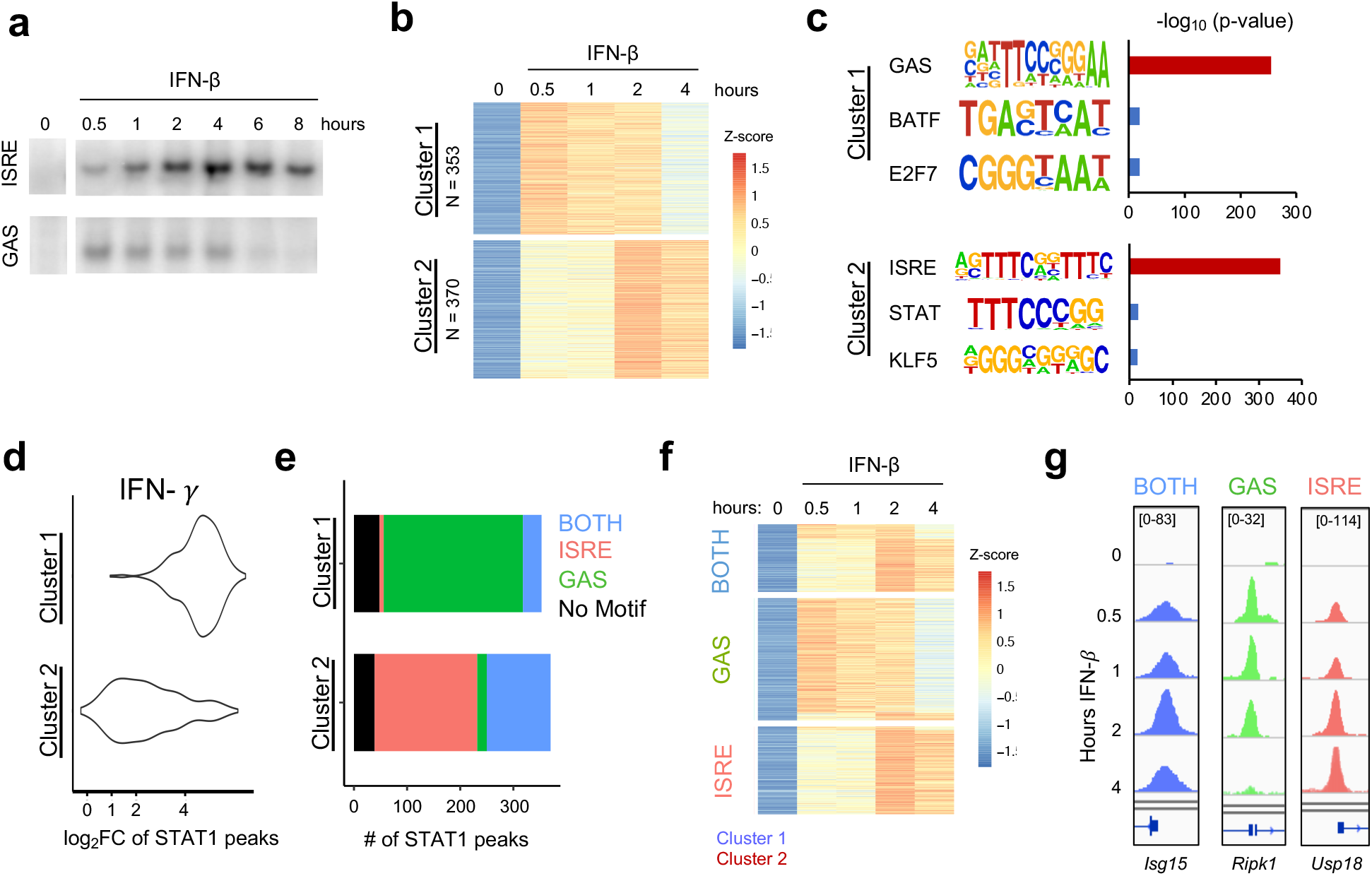
IFN-*β* induced STAT1 binds to GAS and ISRE motifs. **(a)** EMSA of ISRE and GAS binding in MLE12 cells treated with IFN-β (100 U/mL). Data are representative of >5 independent experiments. **(b)** STAT1 ChIP-seq heatmap of inducible STAT1 peaks (FDR < 0.01 for peak calling & > 2-fold induction in at least two time points) after IFN-β (10U/mL) stimulation. Peaks were clustered using k-means clustering. **(c)** Top three hits from *de novo* motif analysis on STAT1 peaks in Cluster 1 *vs*. 2 as defined in (b). **(d)** IFN-γ (100 ng/ml) induced STAT1 ChIP-seq signal comparing locations in Cluster 1 *vs*. 2. **(e)** Stacked bar graph of number of peaks per Cluster that contain ISRE, GAS, or BOTH motifs by searching the HOMER database (see Figure S1). **(f)** Re-clustered STAT1 ChIP-seq heatmap based on motif categories. **(g)** Genome browser tracks of representative promoter-bound STAT1 ChIP-seq peaks from BOTH (*Isg15*), GAS (*Ripk1*), and ISRE (*Usp18*) categories.

**Figure 2:**
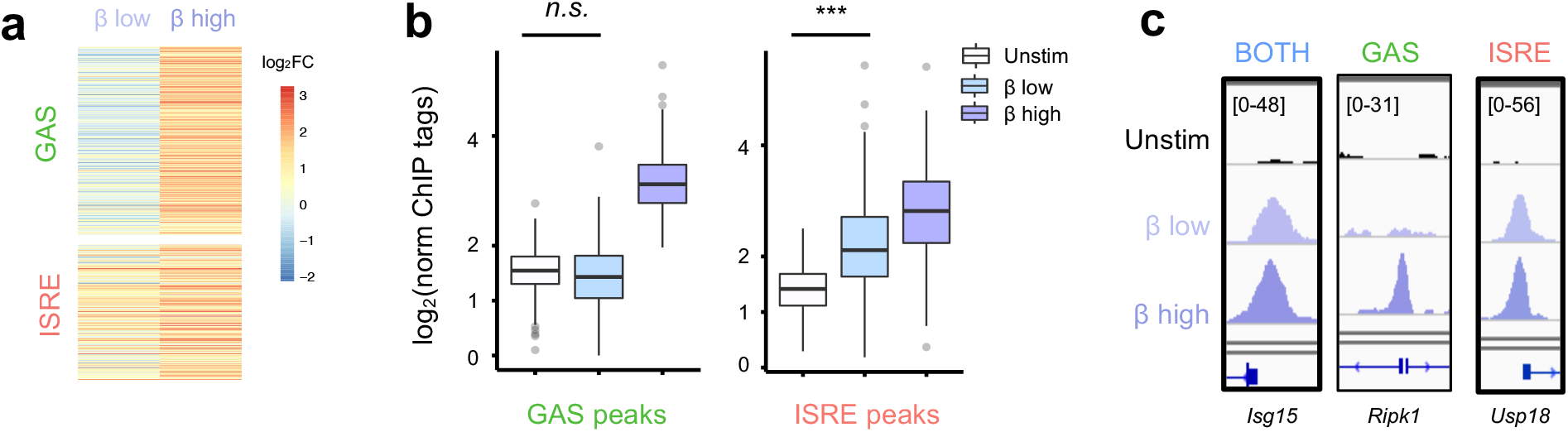
Low-dose IFN-β preferentially activates ISGF3 over GAF. **(a)** Heatmap of STAT1 ChIP-seq with two doses of IFN-! (1 vs 10 U/mL) for 1h, showing GAS and ISRE clusters identified in Fig.1f. **(b)** Boxplot of STAT1 ChIP-seq signals for GAS cluster peaks in (b). *n.s*. = not significant, *** = *p* < 2.2 e-16 by one-way Wilcox ranked sum test. **(c)** Genome browser tracks of representative promoter-bound STAT1 ChIP-seq peaks from “Both” (*Isg15*), GAS (*Ripk1*), and ISRE (*Usp18*) clusters across two doses of IFN-!.

**Figure 3:**
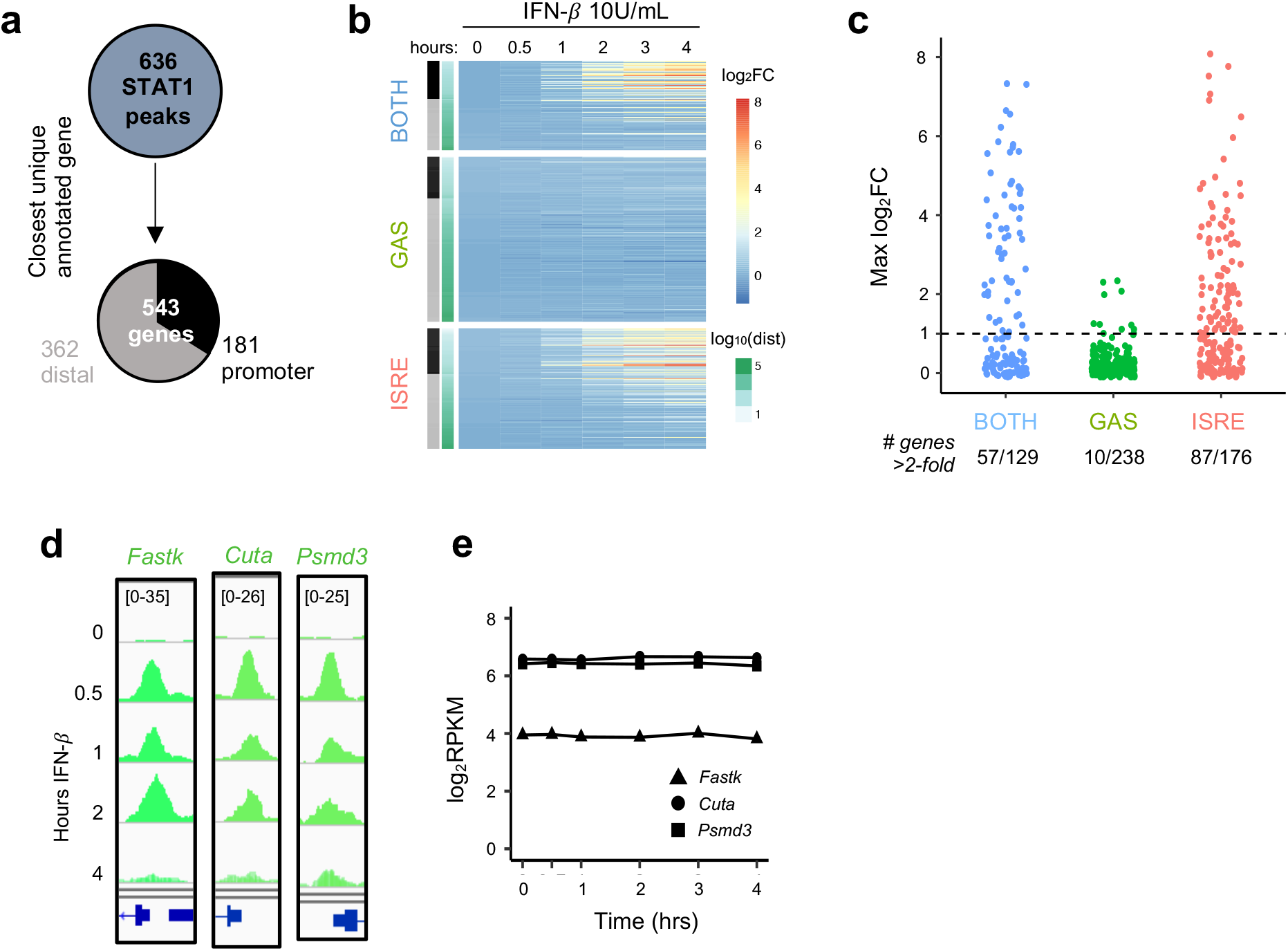
GAF does not induce expression of nearby genes. **(a)** Schematic for linking peaks to genes. Promoter defined as −1000 to +100 bp from TSS. **(b)** Heat map of log2 foldchange of genes linked to STAT1 peaks, clustered by STAT1 peak motifs. Genes are ordered by distance to STAT1 peak. **(c)** Dot plot of genes in each category showing number of genes above 2-fold induction threshold. **(d)** Genome browser tracks of representative STAT1 peaks with GAS motifs at promoters of three genes. **(e)** Gene expression response to IFN-*β* (10 U/ml) for same genes as in (d).

**Figure 4:**
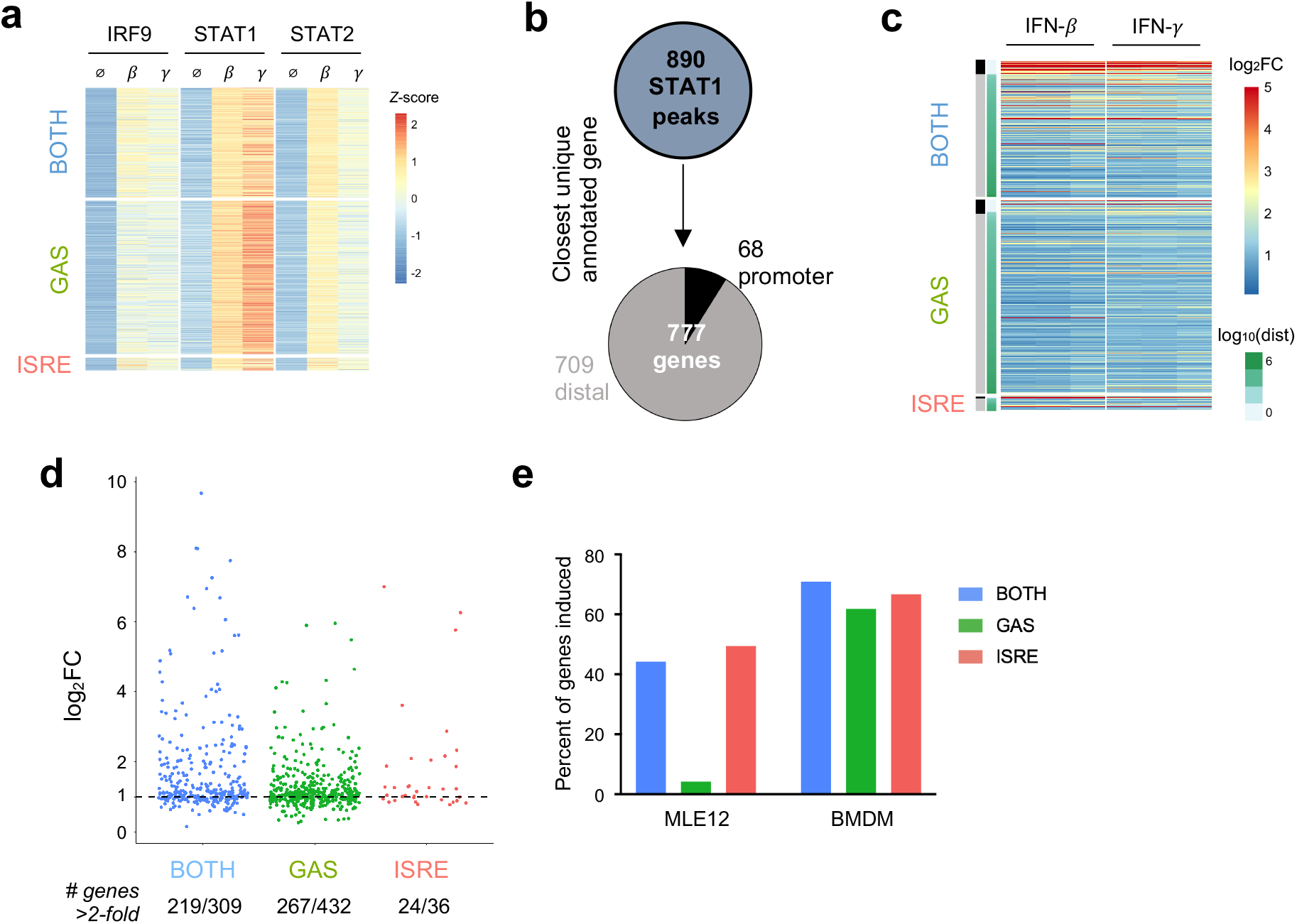
In macrophages, GAF does induce expression of nearby genes. **(a)** Heat map of ChIP-seq data in macrophages (*21*), categorized by presence of GAS or ISRE motif. **(b)** Schematic for linking peaks to genes in macrophages. Promoter defined as −1000 to +100 bp from TSS. **(c)** Heat map of log2 fold-change of macrophage genes linked to macrophage STAT1 peaks. Genes are ordered by distance to STAT1 peak. **(d)** Dot plot of genes in each category showing number of macrophage genes above 2-fold induction threshold. **(e)** Bar graph of the percentage of genes near motif-categorized peaks that are induced upon IFN-*β* stimulation, comparing MLE12 cells *vs*. BMDMs.

**Figure 5:**
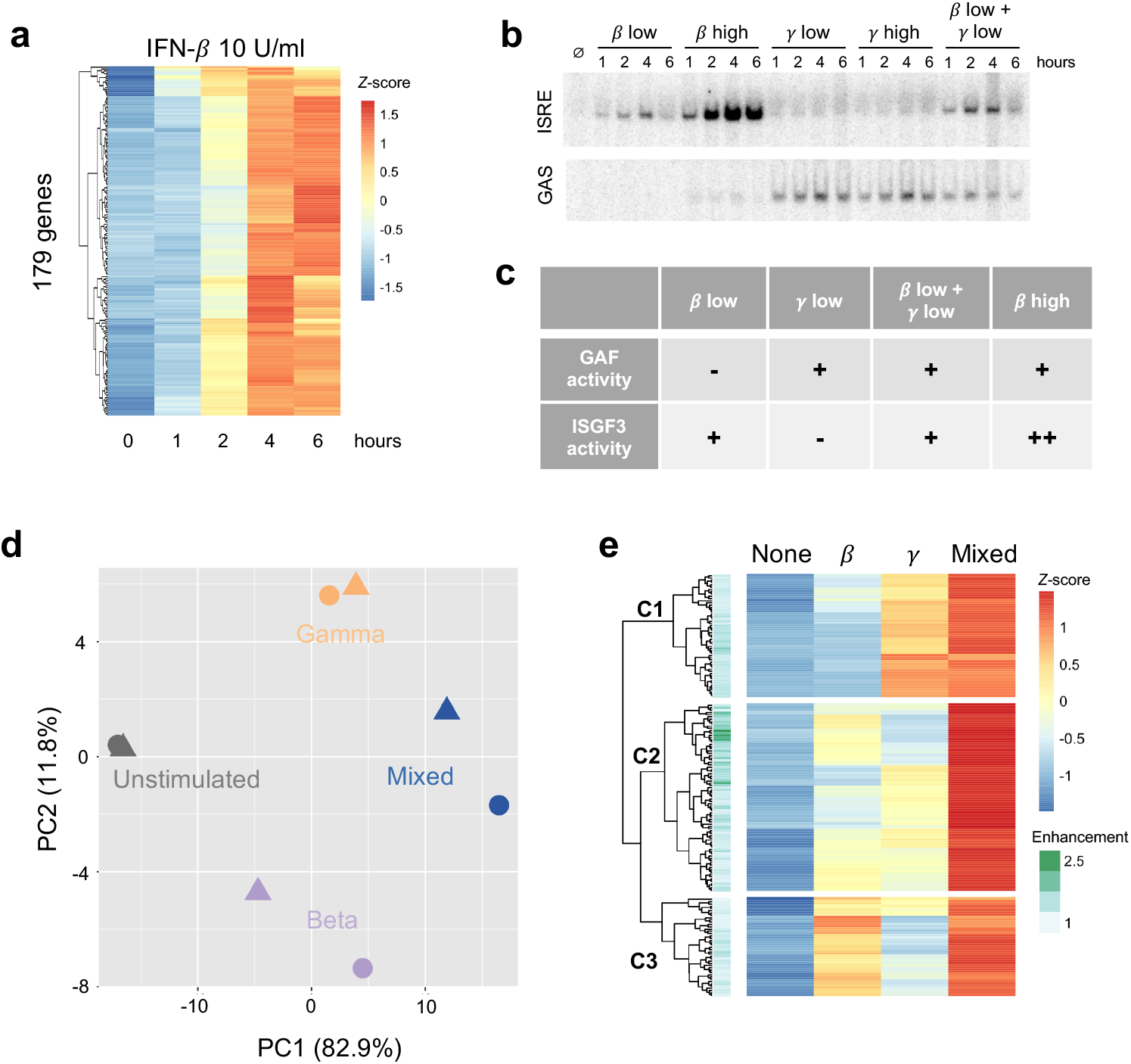
Combined activation of GAF and ISGF3 enhances expression of ISGs in respiratory epithelial cells. **(a)** Heat map of 179 MLE12 ISGs as defined by induction with 10 U/ml IFN-*β* using thresholds of FDR<0.5 and fold-change>2. **(b)** EMSA of ISRE and GAS binding in response to IFN-*β* (1 *vs*. 100 U/mll), IFN-*γ* (1 *vs*. 100 ng/ml), or a mixture of the low doses. Representative gel from five replicates. **(c)** Table of relative ISGF3 and GAF activation strengths in response to different doses of IFN-*β* and IFN-*γ*, based on EMSA data. **(d)** Principal component analysis of ISGs in response to IFN-*β* low dose (1 U/ml), IFN-*γ* low dose (1 ng/ml), or IFN-*β* + IFN-*γ* low doses (“Mixed”). Four-hour stimulation, two replicates. **(e)** Heat map of ISGs showing response to IFN-*β* low-dose, IFN-*γ* low dose, and Mixed. First three levels of hierarchical clustering are highlighted. Row annotations indicate Enhancement score, defined as RPKMmixed / (RPKM_beta_ + RPKM_gamma_)

### BMDM data

Raw ChIP-seq and RNA-seq data (*21*) were downloaded from the NCBI Sequence Read Archive under accession numbers SRP149943 (STAT1, STAT2, and IRF9 ChIP-seq) and SRP149944 (RNA-seq). Data was downloaded as .fastq files and processed in the same manner as data generated in our lab.

### ChlP-seq analysis

The low quality 3’ends of reads were trimmed (cutoff q=30), and remaining adapter sequences were removed using cutadapt (*26*). Reads were aligned using bowtie2 (*27*) using −non-deterministic and −very-sensitive parameters. Peaks were called in each sample against input control by MACS2 peak caller (*28*) with FDR threshold of 0.01 with the following parameters: keep-dup 1 --call-summits --nomodel --extsize 200. Peaks per sample were merged to generate a reference peak list. Each bam file was normalized based on its sequencing depth with deepTools (*29*). Sequencing-depth normalized reads were counted within every peak by multiBigWigSummary command from deepTools to generate a count table. Inducible peaks were identified using a threshold of two-fold induction from the basal condition in at least two timepoints.

One replicate of MLE12 IFN-β stimulation at 0.5 hour had particularly high signal to noise ratio, resulting in 10-fold higher number of peaks called compared to other samples. We included this sample in the identification of possible STAT1 peaks but excluded it for finding inducible peaks, thereby maximizing the number of peaks considered but stringently defining the inducible peaks. K-means clustering analysis was performed on scaled log_2_ normalized ChIP-seq tags. Silhouette and elbow plots were examined to identify the best k of 2 from the data. *De novo* motif analysis was performed within peak regions using the findMotifsGenome function in the HOMER suite (*30*). Heatmaps were generated using the pheatmap package from R (*31*). The April 2019 version of Ensembl database was used to extract transcription start site information and to annotate peaks relative to genes. Bigwig files were generated with the multiBamSummary function in deeptools, and tracks were visualized in IGV (*32*).

### ChIP-seq Motif Clustering

Categorization of STAT1 peaks based on motifs was done by searching the HOMER database (*30*) for position weighted matrix files of STAT/GAS and ISRE/IRF motifs. ISRE/IRF motifs included motifs for T1ISRE, ISRE, IRF2, IRF1, IRF3, and IRF8. STAT/GAS motifs included motifs for STAT1, STAT3+IL-21, STAT3, STAT4, and STAT5 (Fig. S1b). A variant GAS motif in the promoter of *Irf1* (GATTTCCCCGAATG) known to be bound by STAT1 (*33*) was also included in the search as a GAS motif. The annotatePeaks function in HOMER was used to scan for the presence of motifs within inducible STAT1 peaks. A peak that contained at least one kind of STAT/GAS motif was classified as a GAS peak, and a peak that contained at least one kind of ISRE/IRF motif was classified as an ISRE peak. A peak that contained motifs from both STAT/GAS and ISRE/IRF categories was classified as a BOTH peak, while a peak that did not contain either motif was classified as a No motif peak.

### RNA-seq analysis

Reads were trimmed, filtered, and deduplicated in the same manner as ChIP-seq data. Processed reads were aligned to mm10 genome using STAR (*34*), and count tables were generated by the featureCount function in deepTools (*35*). Counts were normalized using TMM-normalization method and RPKM values were generated using edgeR (*36*). Genes below an expression threshold of 2 CPM in all samples were excluded from downstream analysis. IFN-*β* inducible genes (Fig. 3a) were identified by FDR < 0.05 and fold-change > 2 compared to unstimulated. For foldchange and Enhancement score calculations a pseudocount of 1 RPKM was added. Gene ontology analysis was performed using the Database for Annotation, Visualization and Integrated Discovery (*37*). The data were visualized using pheatmap and ggplot2 packages (*31, 38*).

### Electrophoretic Mobility Shift Assay (EMSA)

After stimulation with IFNs, cells were scraped in cold PBS + 1 mM EDTA, collected by centrifugation, and resuspended in Cytoplasmic Extract Buffer (10 mM HEPES-KOH pH 7.9, 60 mM KCl, 1 mM EDTA, 0.5% NP40, 1 mM DTT, and 1 mM PMSF). After incubating on ice for two minutes, cells were vortexed for 30 seconds, and nuclei were collected by centrifugation and resuspended in Nuclear Extract Buffer (250 mM Tris pH 7.5, 60 mM KCl, 1 mM EDTA, 1 mM DTT, and 1 mM PMSF). The nuclei were snap frozen and stored in −80 °C before use.

Nuclear extracts were prepared by disrupting the nuclear membrane with three freeze-thaw cycles of 30 seconds each using 37 °C water bath and dry ice, followed by centrifugation at max speed for 15 minutes. The resulting supernatants containing nuclear proteins were then incubated for 15 mins with P^32^-labeled oligonucleotide probes for 15 minutes in binding buffer (10 mM Tris pH 7.5, 50 mM NaCl, 10% glycerol, 1% NP40, 1 mM EDTA, 0.1 mg/mL polydeoxyinosinic-deoxycytidylic acid). The following probes were used for detecting ISGF3 and GAF DNA-binding activity: ISRE probe (GATCCTCGGGAAAGGGAAACCTAAACTGAAGCC) and GAS probe (TACAACAGCCTGATTTCCCCGAAATGACGC). The reaction mixtures were run on a 5% acrylamide (30:0.8) gel with 5% glycerol and TGE buffer (24.8 mM Tris, 190 mM glycine, 1 mM EDTA) at 200V for 1 hour and 45 mins. The gels were dried and imaged on a Sapphire Biomolecular imager in phosphor mode (Azure Biosystems, Dublin, CA).

### Cycloheximide and qPCR

For cycloheximide experiments, cells were incubated for 20 minutes with media containing 10 *μ*g/ml cycloheximide prior to stimulation with IFN. The following *Cxcl10* primer sequences were used for qPCR: AGTGCTGCCGTCATTTTCTG (forward), ATTCTCACTGGCCCGTCAT (reverse). qPCR reactions were run with Sso Advanced Universal SYBR Green Supermix on a CFX384 RT-qPCR system (BioRad, Hercules, CA).

## Results

### IFN-*β* induced STAT1 binds to GAS and ISRE motifs

To characterize the TFs activated by IFN-β in MLE-12 cells, we performed electrophoretic mobility shift assays (EMSAs) with ISRE and GAS probes. Robust protein binding to both ISRE and GAS probes was detected in response to IFN-β (10 U/ml), indicating inducible activity of ISGF3 and GAF, respectively (Fig. 1a). The GAS signal reached a maximum within one hour post stimulation, while the ISRE signal gradually increased and peaked at four hours.

To examine ISGF3 and GAF binding events genome-wide, we performed STAT1 ChIP-seq at various time points after IFN-β stimulation. A total of 723 inducible STAT1 peaks were identified. K-means clustering revealed two clusters of peaks with distinctive kinetics (Fig. 1b). Cluster 1 contained 353 peaks that were induced most strongly at half hour and continued until two hours after stimulation. Cluster 2 contained 370 peaks that were most strongly induced at two and four hours after stimulation.

As STAT1 is a member of both ISGF3 and GAF, we performed *de novo* motif analysis on the genomic regions in each cluster to infer the identities of STAT1-containing complexes. The peaks in Cluster 1 were highly enriched for a GAS-like motif (*p* = 1e-255), while the peaks in Cluster 2 were highly enriched for an ISRE-like motif (*p* = 1e-350) (Fig. 1c). This suggested that STAT1 peaks in Cluster 1 represent GAF binding events while Cluster 2 peaks represent ISGF3 binding events. To corroborate the result of our motif analysis, we performed STAT1 ChIP-seq in response to IFN-γ stimulation, which strongly activates GAF. As expected, peaks in Cluster 1 showed greater IFN-γ-inducible STAT1 binding than peaks in Cluster 2 (Fig. 1d). These results confirmed the idea that peaks in Cluster 1 represent GAF binding and Cluster 2 represent ISGF3 binding.

We also performed STAT1 ChIP-seq with and without cycloheximide (CHX) to investigate whether GAF binding is dependent on IFN-β-induced *de novo* protein synthesis of secondary signals such as IFN-γ. We found that the addition of cycloheximide did not reduce the ChIP signal in either Cluster 1 or Cluster 2 (Fig. S1a), indicating that IFN-*β* signaling directly activates GAF without the need for protein synthesis.

To complement the unbiased clustering approach, we categorized peaks based on known GAS and ISRE binding motifs. We comprehensively scanned the HOMER database (*30*) for STAT/GAS and ISRE/IRF motifs under the 723 peaks (Fig. S1b). We found that for Cluster 1, 262 out of 353 peaks (74%) had only STAT/GAS motifs, 35 peaks (10%) had both STAT/GAS and ISRE/IRF motifs (BOTH motifs), and 8 peaks (2%) had only ISRE/IRF motifs (Fig. 1e). For Cluster 2, 193 out of 370 peaks (52%) had only ISRE/IRF motifs, 120 peaks (32%) had BOTH motifs, and 18 peaks (5%) had only STAT/GAS motifs (Fig. 1e). In sum, this motif classification analysis resulted in 280 peaks that had only STAT/GAS motifs (categorized as GAS peaks), 155 peaks that had both STAT/GAS and ISRE/IRF motifs (BOTH peaks), and 201 peaks that had only ISRE/IRF motifs (ISRE peaks). A re-clustered heatmap using these categories (Fig. 1f) recapitulated the distinct temporal patterns of STAT1 binding seen by K-means clustering. As examples, peaks from BOTH, GAS, and ISRE motif categories were found in the promoters of *Isg15, Ripk1*, and *Usp18* respectively (Fig. 1g). Genome-wide, the peaks with GAS motifs were on average induced more weakly than peaks with BOTH or ISRE motifs (Fig. S1c). Eighty-seven peaks had neither canonical GAS nor ISRE motifs (Fig. 1e), which may represent STAT1 recruited to chromatin through protein-protein interactions without making direct DNA contacts. Therefore, these peaks were removed from subsequent analyses.

### Low-dose IFN-*β* activates ISGF3 but not GAF

Having defined that IFN-*β* activates GAF genome-wide, we wondered if IFN-*β* dose affected whether GAF is activated. We performed STAT1 ChIP-seq with low (1 U/ml) and high (10 U/ml) doses of IFN-β and observed that at the low IFN-*β* dose, ISRE-containing regions showed inducible STAT1 binding (*p* < 2.2 e-16), yet no significant binding was found at GAS-containing regions (Fig. 2,b) as defined by the motif categorization in Figure 1. For example, the GAS-containing peak at the *Ripk1* promoter is bound in a dose-sensitive manner (Fig. 2c), while peaks at the *Isg15* and *Usp18* promoters that possess BOTH and ISRE motifs, respectively, are induced to a similar magnitude by low and high dose IFN-β. This demonstrated that low-dose IFN-β induces STAT1 binding to ISRE but not GAS sites, suggesting that GAF may act as a molecular switch to distinguish low from high doses of IFN-*β*.

### IFN-*β*-induced GAF does not induce expression of nearby genes

We next interrogated the function of ISGF3 and GAF binding events by linking the 636 inducible STAT1 ChIP-seq peaks identified in Figure 1 (belonging to BOTH, GAS, and ISRE clusters) to their closest expressed genes (Fig. 3a). This yielded 543 uniquely linked genes, of which 181 had promoter STAT1 peaks (−1000bp to + 100bp) and 362 had distal STAT1 peaks. Genes linked to peaks in the BOTH and ISRE clusters showed functional enrichment for defense response to virus and cellular response to IFN-β, confirming that these are known ISGs. Interestingly, however, the 238 genes linked by proximity to GAS peaks were not enriched for any ontology terms (Fig. S2a).

We then examined the expression of the 543 linked genes by performing RNA-seq over a four-hour time course after stimulation with IFN-*β* (10 U/ml). Clustering the genes by the motif categories of their linked STAT1 ChIP-seq peaks (Fig. 3b), we found that genes linked to the BOTH and ISRE clusters were induced upon IFN-*β* stimulation. The magnitude of induction was strongest for genes with promoter STAT1 peaks, while distally linked genes were less likely to be induced. This may be due to the fact that enhancers do not necessarily regulate their closest genes, and the linkage between peak and gene becomes increasingly unreliable with increased genomic distance (*39, 40*).

In striking contrast to the BOTH and ISRE clusters, we found very little induction amongst genes linked to STAT1 peaks with only GAS motifs, even when the binding events were in their promoters (Fig. 3b, GAS cluster). Only 4% of genes (10 out of 238) in the GAS cluster were induced more than two-fold by IFN-*β*, compared to 44% and 49% of genes in the BOTH and ISRE clusters, respectively (Fig. 3c). For example, *Fastk, Cuta*, and *Psmd3* all have prominent IFN-*β*-inducible STAT1 peaks in their promoters that contain GAS motifs (Fig. 3d). However, these genes displayed no IFN-*β*-inducible change in expression level (Fig. 3e).

Genes linked to GAS peaks were also not inducible by higher doses of IFN-β and IFN-*γ* Even at 100 U/ml IFN-*β* and 100 ng/ml IFN-*γ*, only eight and five out of 238 genes, respectively, were induced greater than two-fold (Fig. S2b,c). One of the few GAS-linked genes that was induced by both IFN-*β* and IFN-*γ* was *Irf1*, a known IFN-*γ* target gene that has been implicated as a key factor in amplifying IFN responses (*41*). *Irf1* may thus be an important exception to the general observation that GAF binding does not induce expression of nearby genes.

### In macrophages, GAF does induce expression of nearby genes

Given that IFN-*γ* has a well-described role in macrophage function (*20*), we wondered if GAF was similarly inactive in a macrophage context. Using publicly available datasets of STAT1, STAT2, and IRF9 ChIP-seq and RNA-seq in bone marrow-derived macrophages (BMDMs) (*21*), we performed an identical analysis to what was done in MLE12 cells. A total of 890 STAT1 peaks were inducible by 90-minute stimulations with either IFN-*β* (200 U/ml) or IFN-*γ* (10 ng/ml). These were categorized into 469 peaks containing GAS motifs only, 38 peaks containing ISRE motifs only, and 335 peaks containing BOTH motifs (Fig. 4a). The low number of ISRE-containing STAT1 peaks was due partly to the relatively weak STAT1 signal and also consistent with prior observations that many ISGF3 binding events in this system are STAT1-independent (*21*). To overcome this limitation, we used an alternative classification method and categorized BMDM ChIP-seq peaks by patterns of TF binding: locations with STAT1 peaks and no IRF9 peaks were considered GAF binding events, and locations with IRF9 peaks in WT cells and no STAT1 or STAT2 peaks in *Irf9*^-/-^ cells were considered ISGF3 binding events. Using this approach, we identified 1281 ISGF3 and 677 GAF binding events (Fig. S3a, S3b).

We then linked peaks to their nearest expressed genes to examine IFN-responsive gene expression for both categorization methods. Using the motif-based categorization of STAT1 peaks, we identified 777 uniquely linked genes (Fig. 4b). In contrast to our observation in MLE12 cells, here we found that genes linked to GAS peaks were induced in a similar manner as genes linked to ISRE or BOTH peaks (Fig. 4c), with similar patterns of induction by IFN-*β* or IFN-*γ* stimulation. We found that 61.8% of genes linked to GAS peaks, 66.7% of genes linked to ISRE peaks, and 70.9% of genes linked to BOTH peaks were induced greater than two-fold. The alternative method of classifying peaks by TF binding reproduced this result, with 59.1% of genes linked to GAF binding and 61.6% of genes linked to ISGF3 induced greater than two-fold (Fig. S3c).

These findings indicated that GAF is transcriptionally active in macrophages but not in epithelial cells (Fig. 4e), likely due to the presence of collaborating TFs that are expressed in macrophages (*42, 43*). Indeed, promoters of genes linked to GAS peaks were enriched 1.9-fold (*p* = 10e-7) for motifs of PU.1, a macrophage-specific transcription factor that collaborates with STAT1 in gene expression (*43, 44*).

### Combined activation of GAF and ISGF3 enhances expression of ISGs in respiratory epithelial cells

Given the surprising finding in epithelial cells that IFN-*β* activates substantial GAF binding with no direct transcriptional activity, we hypothesized that GAF might play an indirect role in the regulation of ISGs. To address this hypothesis, we first defined ISGs using our RNA-seq data from MEL12 cells stimulated with high dose IFN-*β* (10 U/ml). We identified 179 ISGs (Fig. 5a) by stringent statistical thresholds (FDR < 0.05 and fold-change > 2) and used this set of genes for subsequent analysis. To study the effects of GAF on ISG expression, we leveraged the fact that low-dose IFN-β activates ISGF3 but not GAF (Fig. 2), while low-dose IFN-γ activates GAF but not ISGF3 (Fig. 5b). We stimulated cells with low-dose IFN-β, low-dose IFN-γ, or a mixture of the two, resulting in the TF activation patterns show in Fig. 5c. Comparing gene expression in the Mixed condition and the single-stimulus conditions thus allowed for discovery of genes that are co-regulated by ISGF3 and GAF.

We performed RNA-seq in response to IFN-β, IFN-*γ*, and the Mixed condition at four hours after stimulation and examined the expression of the 179 previously-defined ISGs. Principal component analysis showed that two replicates of the Mixed condition fell between IFN-β and IFN-*γ* samples and were placed further away from the unstimulated sample compared to the single stimulus conditions (Fig. 5d). This suggested that the Mixed condition resulted in a gene expression profile that is a hybrid of IFN-β & *γ* profiles and that some ISGs were most strongly induced in the Mixed condition.

Visualizing these ISGs in a hierarchically clustered heatmap provided additional insight into the effects of GAF and ISGF3 (Fig. 5e). We observed that ISGs in the top cluster (C1) were induced by IFN-*γ* alone, and their expression was only mildly enhanced in the Mixed condition. Similarly, genes in the bottom cluster (C3) were induced by IFN-*β* alone and mildly enhanced in the Mixed condition. Thus, C1 and C3 genes demonstrated stimulus-specific patterns of expression, and, interestingly, motif analysis of both clusters revealed enrichment of IRF and ISRE motifs but not GAS motifs in the promoters.

The expression of ISGs in the middle cluster (C2) was of greatest interest. These ISGs were induced weakly in response to either IFN-*β* or IFN-*γ* alone, but strongly enhanced in the Mixed condition. To quantify the effect of mixing IFN-*β* and IFN-*γ*, we calculated an Enhancement score for each gene: Enhancement score = RPKM_mixed_ / (RPKM_β_ + RPKM_*γ*_) (Fig. 5e annotation). This objective score confirmed that many genes in the middle cluster are strongly enhanced in the Mixed condition, suggesting that ISGF3 and GAF may co-regulate the expression of some ISGs.

To investigate the properties of highly enhanced genes, we compared ISGs in the top quartile of Enhancement score (n = 45, Enhancement > 1.156) to ISGs in the bottom quartile (n = 45, Enhancement < 0.774)) (Fig. 6a). To confirm that enhanced gene expression occurred over multiple time points, we performed RNA-seq over a six-hour time course. Genes in the Top quartile indeed displayed a pattern of enhancement across multiple time points, and the degree of inducibility in the Mixed condition approached that seen with high dose IFN-β (Fig. 6b). We also observed that genes in the Top category were induced more robustly than genes in the Bottom category for all conditions (Fig. 6b,c). This was primarily due to the fact that genes in the Top category had much lower basal expression than genes in the Bottom category (Fig. 6c, *p* = 2.66 e-15 for difference between basal RPKMs).

**Figure 6:**
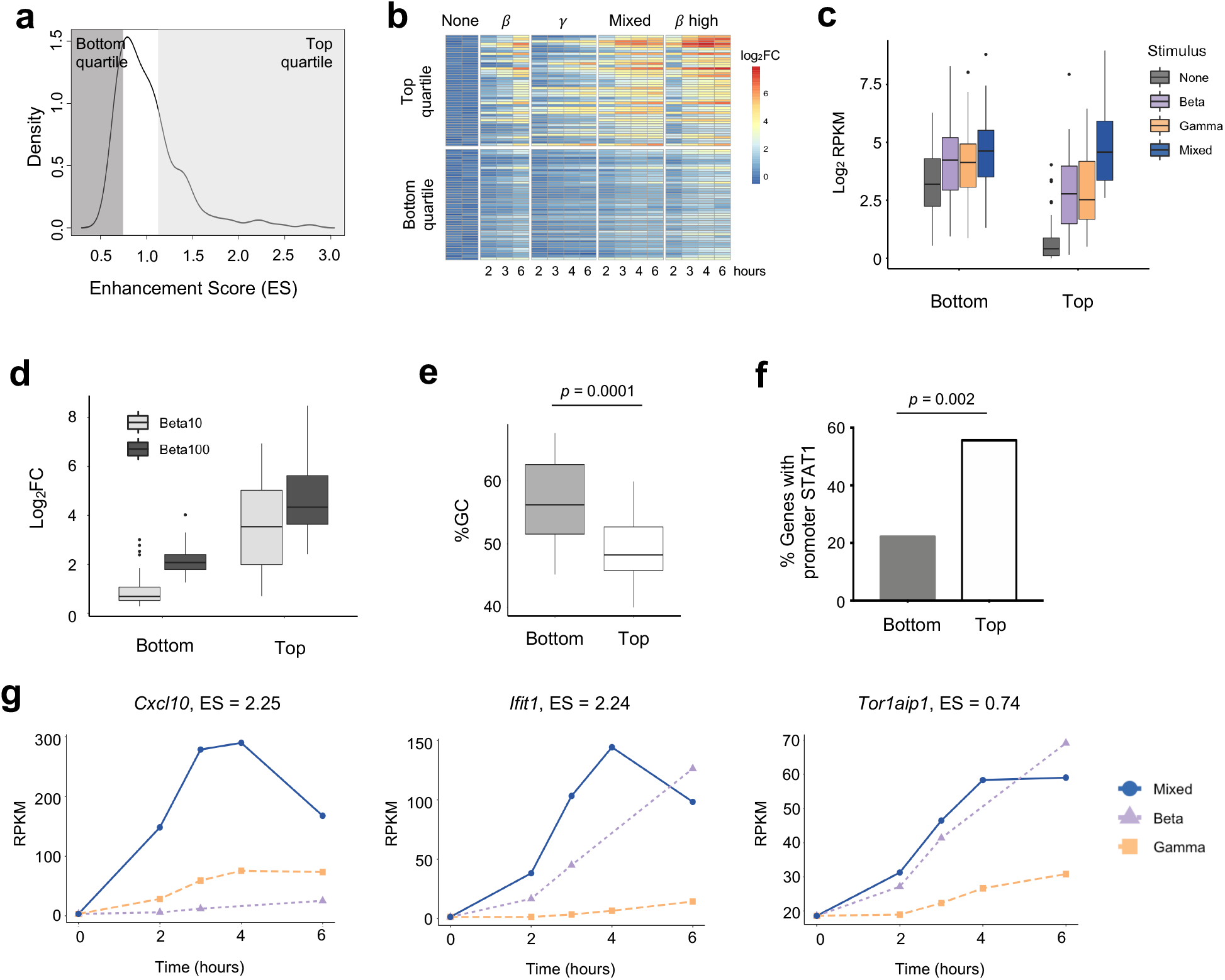
Properties of strongly enhanced ISGs. **(a)** Density plot showing distribution of Enhancement Scores (ES) across 179 ISGs. Genes in the Top quartile (ES > 1.156) and Bottom quartile (ES < 0.774) are shaded (n=45 genes each). **(b)** Heat map of top and bottom quartile genes in a time course of stimulation of IFN-*β* low, IFN-*γ* high, Mixed, and IFN-*β* high (10 U/ml). **(c)** Box plots of absolute gene expression in Log_2_RPKM for top and bottom enhancement genes by stimulus. **(d)** Box plot of GC content at TSS (−300bp to +300bp) of top *vs*. bottom enhancement genes. **(e)** Percentage of top *vs*. bottom enhancement genes that have a STAT1 ChIP-seq peak in promoter (−1000bp to +100bp). **(f)** Box plots of inducible expression of top *vs*. bottom enhancement genes in response to high doses of IFN-*β* (10 and 100 U/ml). **(g)** Line plots of representative genes of Top (*Cxcl10, Ifit1*) *vs*. Bottom enhancement score (*Torlaip1*).

We confirmed that genes in the Top category were also substantially more inducible in response to high doses of IFN-β (10 and 100U/mL, Fig. 6d, suggesting that the Enhancement score in our mixing experiment is predictive of gene expression responses at high doses of IFN-β that activate both GAF and ISGF3.

Prior studies have shown that GC content is anti-correlated with nucleosomal stability, and that genes with high GC content in their promoters are constitutively expressed because they do not require nucleosome remodeling for transcriptional activation (*45, 46*). We calculated GC content in promoters (−300bp to +300bp) of Top *vs*. Bottom Enhancement score genes and found that promoters of the Bottom category genes showed significantly higher GC content compared to promoters of the Top category genes (*p* = 0.0001, Fig. 6e). We also found that 25 out of 45 (56%) genes in the Top enhancement quartile had STAT1 ChIP-seq peaks in their promoters (−1000bp to +100bp) at any time point, compared to only 10 out of 45 (22%) genes in the Bottom quartile (*p* = 0.002, Fig. 6f). These findings support the idea that genes most strongly enhanced by the combined activation of ISGF3 and GAF are not constitutively expressed but require inducible TF binding at their promoters to establish a chromatin environment that permits maximal expression.

Genes in the Top quartile of Enhancement score included the pro-inflammatory cytokine *Cxcl10* (Enhancement score = 2.25) and the antiviral effector *Ifit1* (Enhancement score = 2.24), both of which demonstrate low basal expression and greatest inducibility in the Mixed condition (Fig. 6g). In contrast, *Tor1aip1* was representative of genes in the Bottom quartile, with relatively high expression at the basal state (19 RPKM) and modest inducibility that was similar in IFN-*β* alone and the Mixed condition. A complete list of Top and Bottom quartile genes is provided in Table S1. TF motif analysis and gene ontology analysis did not reveal any difference between the two categories. Both Top and Bottom genes contained ISRE motifs in their promoters and were enriched for IFN-related ontology terms such as “Viral defense” and “Immune response” (data not shown).

### Gene-specific mechanisms of GAF-mediated ISG enhancement

If GAF binding does not directly induce expression of nearby genes, yet mixing IFN-*β* and IFN-*γ* enhances expression of ISGs, we hypothesized that there must be indirect mechanisms by which GAF cooperates with ISGF3. One possible mechanism is that GAF and ISGF3 may work cooperatively at the promoters of enhanced genes (Fig. 7a), where GAF may either recruit general TFs or increase chromatin accessibility to facilitate ISGF3 binding. Consistent with this, genes with promoter STAT1 peaks containing BOTH motifs had higher enhancement scores compared to genes with promoter STAT1 peaks containing only ISRE motifs (Fig. 7b), implying that combinatorial binding of GAF and ISGF3 at a BOTH peak may enhance gene expression. For example, *Mx2* had two distinct STAT1 peaks in its promoter, one of which had BOTH motifs, and it had an enhancement score of 1.23 (Fig. 7c).

**Figure 7:**
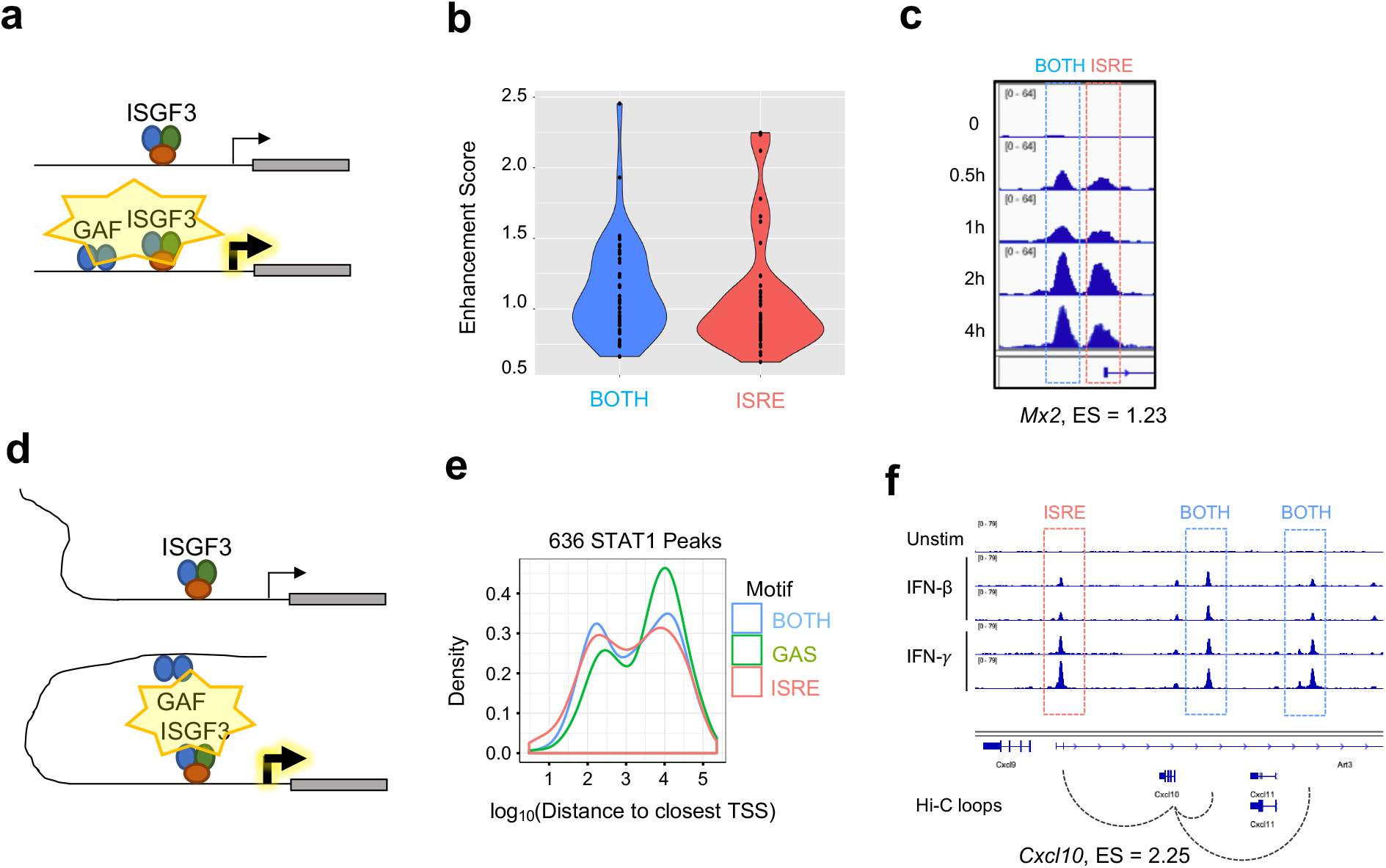
Gene-specific mechanisms of GAF enhancement. **(a)** Schematic of GAF and ISGF3 colocalized at promoters to enhance ISG expression. **(b)** Violin plots of Enhancement scores for genes with STAT1 promoter peaks containing BOTH motifs (GAS+ISRE) *vs*. ISRE motif alone. **(c)** STAT1 ChIP-seq tracks at *Mx2* promoter which has a BOTH peak in the proximal promoter as well as an ISRE peak at the TSS. **(d)** Schematic of distal GAF binding looping to promoters of ISGs to enhance ISG expression. **(e)** Density plot of distance from all STAT1 peaks to nearest TSS, separated by peak motif category. **(f)** Genome browser tracks of STAT1 ChIP-seq data and Hi-C data (*47*) at *Cxcl10* locus demonstrating contact of two BOTH peaks with the TSS.

A second possible mechanism is that GAF may bind to distal enhancers of ISGs (Fig. 7d). GAS peaks were distributed further away from annotated genes than BOTH or ISRE peaks (Fig. 7e), and the nearest genes to GAS were not enriched for IFN-related ontology terms (Fig. S2a). This suggested that GAF binds predominantly in intergenic regions functioning as distal enhancers rather than promoters. To explore this hypothesis, we examined promoter-capture Hi-C (Pc-HiC) data from mouse embryonic stem cells (*47*). Genome-wide analysis of this Pc-HiC data did not identify significant differences between Top and Bottom Enhancement score genes, likely due to the fact that enhancers are highly cell type specific (*48*). However, we found that the promoter of *Cxcl10*, a gene with high enhancement score, makes statistically significant Hi-C contacts with three neighboring STAT1 peaks, of which two contain BOTH motifs and one contains an ISRE motif (Fig. 7f).

A third possible mechanism is that GAF directly induces expression of a key signaling protein that amplifies the IFN response. This could include signaling components of the ISGF3 pathway, or secondary TFs such as IRF1 (*22*), which was one of the eight inducible genes with GAS peak in its promoter (Fig. S4a-c). GAF-induced IRF1 may then cooperate with ISGF3 at promoters by binding to the same ISRE motifs (*49*). To examine the possibility of GAF enhancing ISG expression through a secondary mediator like IRF1, we treated MLE12 cells with cycloheximide (CHX) and stimulated with low dose IFN-*β*, low dose IFN-*γ*, and both IFNs mixed as before. We found that the enhancement of *Cxcl10* was preserved even in CHX treated cells (Fig. S4d), indicating that enhancement of *Cxcl10* does not require synthesis of a secondary signaling protein.

## Discussion

Coordinated activation of both ISGF3 and GAF in response to IFN-*β* stimulation has been previously observed, but the function of GAF in this context has remained unclear. We found in epithelial cells that activation of GAF occurs only at high doses of IFN-*β*, whereas low-dose IFN-β activates only ISGF3 but not GAF. Surprisingly, STAT1 binding to GAS sites was insufficient to induce gene expression in respiratory epithelial cells but not BMDMs. Despite the lack of GAF-mediated transcription, we observed that stimulating epithelial cells with low doses of IFN-β and IFN-*γ* together, which recapitulates the coordinated activation of ISGF3 and GAF by high doses of IFN-β, enhanced ISG expression compared to either stimulus alone. We posit that GAF enhances ISG expression through a variety of gene-specific mechanisms that enhance the activation potential of ISGF3.

Our finding that IFN-*β* activates GAF in a dose-sensitive manner carries important implications for immunity and inflammation. Low levels of tonic IFN-*β* maintain homeostasis in multiple tissues and confer a basal degree of antiviral protection (*50–52*). Our study suggests that in tonic IFN-*β* conditions, the expression of ISGs is restricted to low levels in the absence of GAF. However, in the context of an active infection, IFN-*β* concentration in the microenvironment increases, resulting in activation of GAF, which acts as a molecular switch to enhance expression for a subset of ISGs. These enhanced ISGs include antiviral mediators such as *Ifit1*, as well as pro-inflammatory cytokines such as *Cxcl10* (Fig. 6g). The absence of GAF in tonic IFN-*β* conditions may thus restrict the expression of pro-inflammatory genes that would be harmful in homeostatic conditions. Consistent with this, a recent study demonstrated that IFN-*β* induces pro-inflammatory genes through the STAT1-IRF1 axis in human hepatocyte cell lines (*22*). Taken together, we propose a model where ISGs fall into two classes: those that are GAF-independent and weakly induced even at high IFN-*β* doses, and those that are enhanced by when GAF is activated. Thus the IFN-*β*-GAF axis controls the ouput of a subset of ISGs in a dose-sensitive manner to limit inflammation at homeostasis but promotes both antiviral and pro-inflammatory gene expression in response to pathogens.

A striking conclusion of our study was that STAT1 binding to GAS elements is insufficient to activate gene expression in epithelial cells. Genes linked to GAF binding events were not induced in response to high-doses of either IFN-*β* or IFN-*γ*, even when the binding events were in their promoters (Fig. 3 and Fig. S2). In contrast, using an identical analysis in BMDMs we found that GAF binding is associated with nearby gene expression in macrophages. The likely explanation for this cell type-specificity is the presence of collaborating TFs that are more abundant or basally bound in macrophages, such as IRF1, IRF8, and/or PU.1 (*43*), thus providing a chromatin environment that better facilitates transcription.

Our findings imply that in certain contexts GAF alone is insufficient for transcriptional activation. This has been suggested by others, and there are two plausible reasons why this may be true. First, there are two isoforms of STAT1: full-length STAT1α and a shorter STAT1*β* that lacks the S727 residue required for maximum transcriptional activity (*53–55*). STAT1*β* poorly with transcriptional co-activators such as CBP/p300 and thus has reduced transcriptional activity compared to STAT1α (*56–58*). It has been suggested that STAT1α preferentially forms ISGF3 complexes while the less active STAT1*β* forms GAF complexes (*53*). Second, post-translational modifications are required for STAT1 transactivation. Unphosphorylated STAT1 can participate in transcription as part of the ISGF3 complex (*57, 59*) due to the transactivation activity of STAT2. But as a homodimer, S727 phosphorylation is required for STAT1 transcriptional activity (*57*). Future ChIP experiments using antibodies against putative collaborating TFs or different STAT1 isoforms and phosphorylation states would shed light on these proposed mechanisms.

Despite its inability to directly activate gene expression in epithelial cells, GAF enhances the expression of ISGs. Our results indicate three distinct potential mechanisms. First, GAF may exert almost all of its gene expression effects through the induction of a secondary TF, IRF1. Induced IRF1 may then enhance ISG expression by binding to the same ISRE motifs as ISGF3. The importance of this GAF-IRF1 axis has been previously described (*25*), and our findings extend this model by proposing that IRF1 may in fact be one of the only genes that is a direct target of GAF. In this way, the GAF-IRF1 axis may be analogous to the IRF3-ISGF3 axis, where the secondary TF is a far more potent activator of gene expression than the primary TF, whose principal role is to induce the secondary TF.

While the induction of IRF1 is an important mechanism for GAF-mediated ISG enhancement, it may not apply to every ISG. We propose two additional mechanisms by which GAF may collaborate with ISGF3 to induce maximal gene expression: First, GAF may colocalize with ISGF3 at promoters. Second, distally-bound GAF may interact with promoter-bound ISGF3 through chromosomal looping. Both of these mechanisms imply an indirect role for GAF in modulating the chromatin environment to facilitate ISGF3 binding and transactivation. STAT1 has a well defined epigenetic role downstream of IFN-*γ* in macrophages, priming genes for greater response to other TFs such as NFκB by modulating the chromatin environment (*20, 60–62*). A similar mechanism, albeit on a shorter time scale, may underlie GAF’s enhancement of ISGF3 in response to high IFN-*β* doses. Intriguingly, our ChIP-seq time course data show that GAF activity peaks earlier than ISGF3, suggesting that GAF may bind first, altering chromatin states and facilitating subsequent ISGF3 binding and robust ISG induction.

Our study demonstrates that IFN-*β*-induced GAF collaborates with ISGF3 to enhance expression of ISGs. These results advance our understanding of the regulation of type I IFN signaling while raising a number of questions ripe for further investigation. Whether GAF primarily enhances ISG expression through inducible IRF1 or other mechanisms should be explored further with genetic perturbations of *Irf1* or the GAS site in its promoter. Further inquiry into the epigenetic changes induced by GAF binding could include measurements of chromatin accessibility or chromosome conformation at GAF binding sites. Finally, the physiological role of GAF in the IFN-*β* response should be explored using *in vivo* models of infection in which its coordinated activation is dysregulated.

## Supporting information

Graphical Abstract

## Author Contributions

KK, CW, JB, and QC performed the experiments. KK, MN, CO, and QC analyzed the data. KK and QC wrote the manuscript. AH and QC conceived of the study. AH obtained funding for the study.

## Acknowledgements

The work was funded by NIH grant (to AH) R01AI132835. KK, MN, and CO were funded by the UCLA Bruins in Genomics Undergraduate Research Program. CW was supported by Ruth L. Kirschstein National Research Service Award AI007323 and the Chancellor’s Postdoctoral Fellowship at UCLA. JB was supported by the Irving and Jean Stone Foundation and the Oppenheimer Award of the UCLA Undergraduate Research Fellows Program. QC was supported by the UCLA Department of Medicine Specialty Training and Advanced Research Program.

