## Supplementary material for "High dose IFN-*β* activates GAF to enhance expression of ISGF3 target genes in epithelial cells": Graphical Abstract

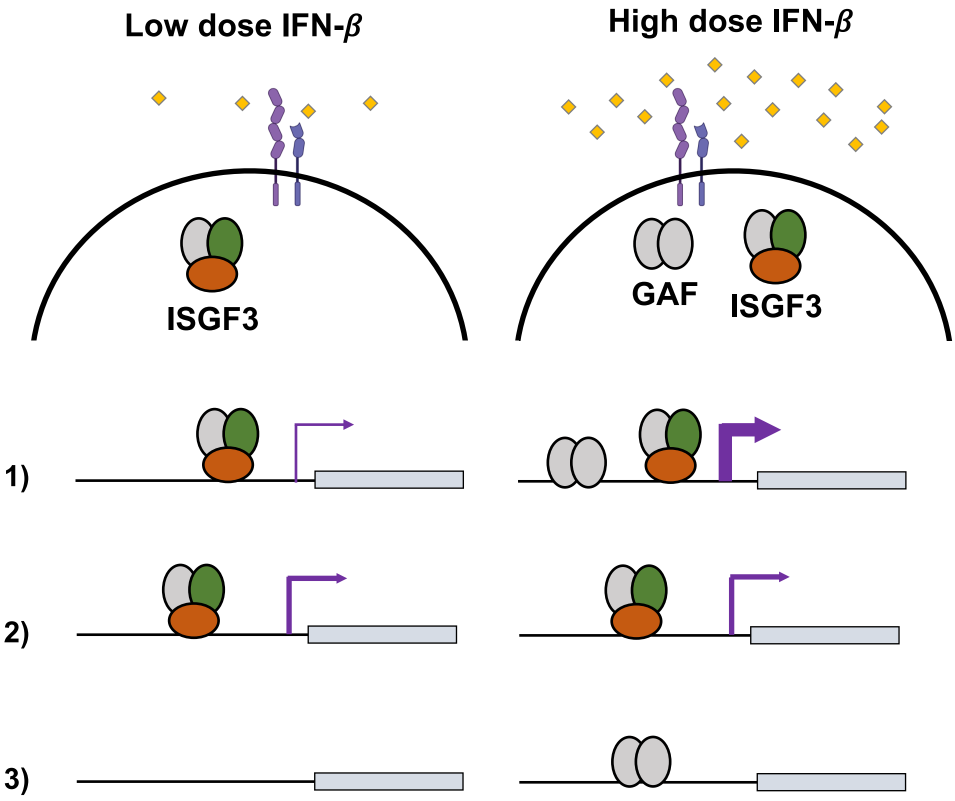


1) Some ISGs are enhanced by GAF at high doses of IFN-$\beta$
2) Other ISGs are not enhanced by GAF
3) GAF binding alone does not induce gene expression
