## Supplemental Figures for "High dose IFN-*β* activates GAF to enhance expression of ISGF3 target genes in epithelial cells"

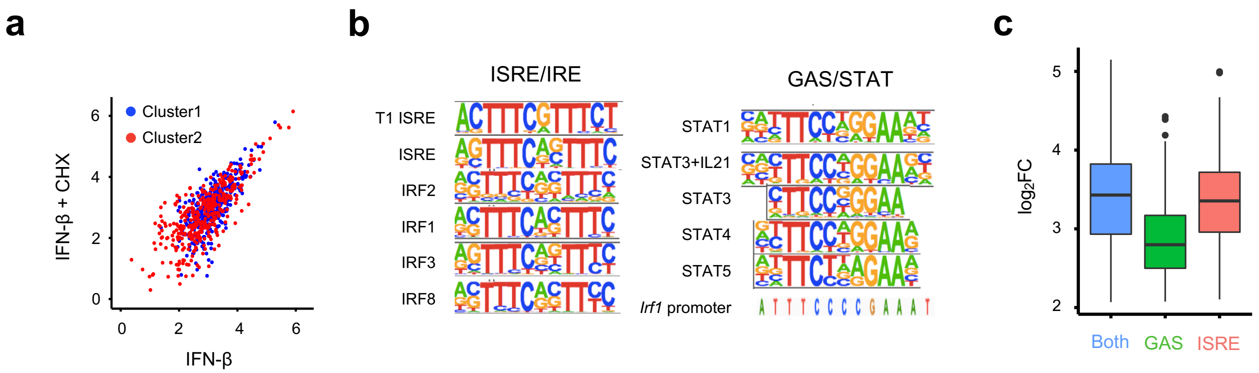


**Figure S1: Supplemental STAT1 ChIP-seq data. (a)** Scatterplot of STAT1 ChIP-seq signal, $\log_2$ of normalized tag counts for 723 peaks, comparing IFN-$\beta$ stimulation (10 U/ml) with and without cycloheximide. Blue dots are Cluster 1 peaks, red dots are Cluster 2 peaks as defined by K-means clustering in Fig 1b. **(b)** JASPAR matrices from the HOMER database used in motif-based classification of STAT1 ChIP-seq peaks. **(c)** Box plots of maximum $\log_2$ fold-change over four-hour time course for STAT1 peaks containing BOTH, GAS, or ISRE motifs.

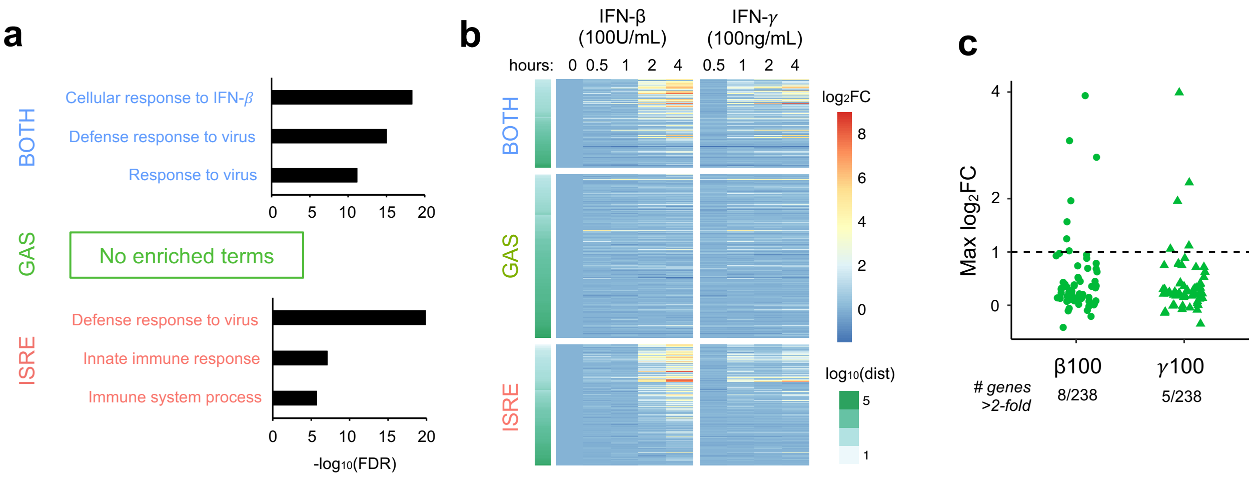


**Figure S2: Supplemental characterization of genes linked to GAS peaks. (a)** Ontology analysis of genes linked to STAT1 peaks containing Both, GAS, or ISRE motifs. **(b)** Heat map of inducible expression ($log_2$ fold-change) for genes linked to STAT1 peaks containing Both, GAS, or ISRE motifs in response to very high doses of IFN-$\beta$ (100 U/ml) or IFN-$\gamma$ (100 ng/ml) in MLE-12 cells. **(c)** Dot plot of $log_2$ fold-change for genes linked to GAS peaks, in response to very high dose IFN-$\beta$ and IFN-$\gamma$ as in (b). Dotted line indicates 2-fold induction threshold.

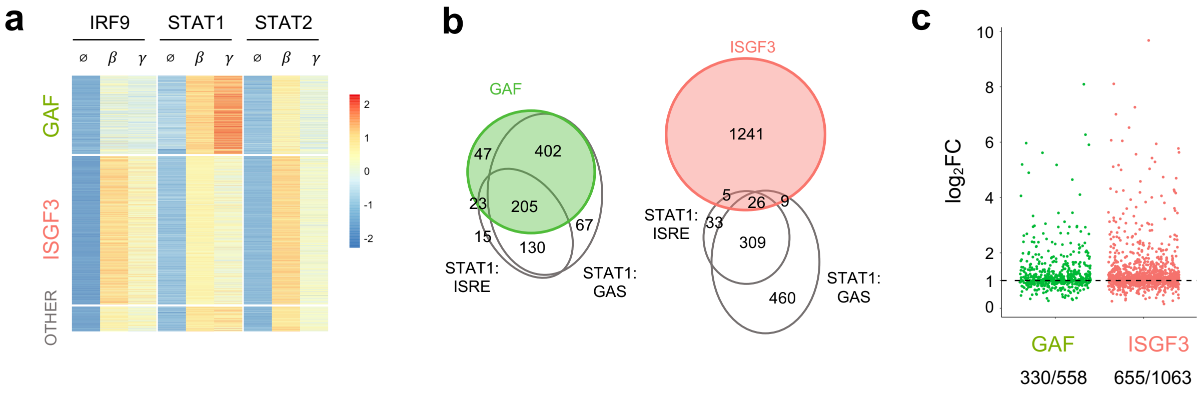


**Figure S3: Categorization of BMDM ChIP-seq peaks by TF binding pattern. (a)** Heat map of ChIP-seq data in macrophages *(21)*, categorized into GAF and ISGF3 peaks by TF binding pattern. **(b)** Venn diagrams comparing peak categorization by TF binding pattern (colored circles, as in Fig. S3a) *vs.* by motifs in STAT1 peaks (gray circles, as in Fig. 4). **(c)** Dot plot of $\log_2$FC expression of genes linked to peaks categorized by TF binding pattern.

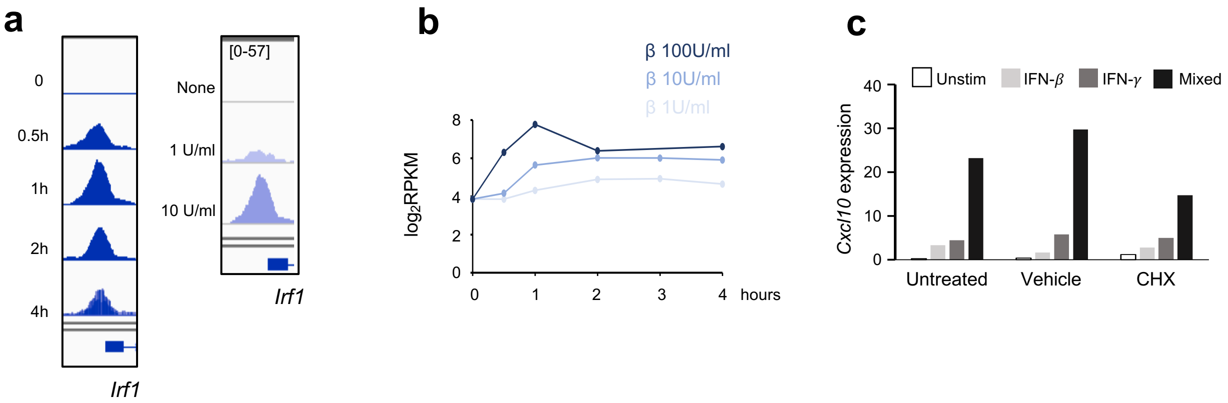


**Figure S4: Role of inducible IRF1. (a)** Genome browser tracks of STAT1 ChIP-seq at *Irf1* promoter. Left, time course with 10 U/ml IFN-$\beta$. Right, low vs high-dose IFN-$\beta$. **(b)** Expression of *Irf1* in response to different doses of IFN-$\beta$. **(c)** qPCR for expression of *Irf1* four hours after stimulation with IFN-$\beta$ (1 U/ml), IFN-$\gamma$ (1 ng/ml), or a mixture of the two, with or without cycloheximide (CHX). Vehicle = DMSO. Expression = percent of GAPDH control.
